# Dynamic plasticity within the EMT spectrum, rather than static mesenchymal traits, drives tumor heterogeneity and metastatic progression of breast cancers

**DOI:** 10.1101/2021.03.17.434993

**Authors:** Meredith S. Brown, Behnaz Abdollahi, Owen M. Wilkins, Priyanka Chakraborty, Nevena B. Ognjenovic, Kristen E. Muller, Mohit Kumar Jolly, Saeed Hassanpour, Diwakar R. Pattabiraman

**Affiliations:** Department of Molecular and Systems Biology, Geisel School of Medicine at Dartmouth, Hanover NH 03755 USA; Department of Biomedical Data Science, Geisel School of Medicine at Dartmouth, Hanover NH 03755 USA; Genomics and Molecular Biology Shared Resource, Norris Cotton Cancer Center, Geisel School of Medicine, Lebanon, New Hampshire, USA; Centre for BioSystems Science and Engineering, Indian Institute of Science, Bengaluru, India 560012; Department of Pathology, Dartmouth-Hitchcock Medical Center, Lebanon, NH 03756 USA

## Abstract

The Epithelial-to-Mesenchymal Transition (EMT) is a developmental cellular program frequently coopted by cancer cells^1^ and is a key contributor to both heterogeneity in solid tumors^2–4^ and later stage chemo-resistance and metastasis^5,6^. Rather than being a switch from an epithelial to a mesenchymal state, increasing evidence points to the existence of intermediate EMT states, wherein cells co-express both epithelial and mesenchymal traits^7–13^. Multiple stable intermediate EMT states possessing unique characteristics exist across the EMT spectrum^7,8,14,15^, contributing to the complex heterogeneity of tumors to promote metastasis^16^. While much work has been carried out identifying and characterizing EMT-inducing transcription factors^17–20^, the transcriptional and epigenetic networks responsible for the stability and maintenance of the midpoints along the EMT spectrum are poorly defined. In addition, there are currently no approaches to identifying and quantifying intermediate EMT subpopulations within patient tumors to evaluate their prognostic significance. Using clonally isolated derivatives of the SUM149PT breast cancer cell line, we systematically interrogate how each EMT state independently contributes to heterogeneity and influences metastatic progression, uncovering the role of RUNX2 in stabilizing certain intermediate states. Using SUM149PT-derived tumors as a training set, we develop an entropy-based model to quantify phenotypic heterogeneity and EMT status. Remarkably, the majority of cell states captured in the SUM149PT model are represented in a selection of patient tumors, laying the foundation for quantification of epithelial-mesenchymal heterogeneity and understanding the role of the intermediate EMT state in tumor progression.

---

We derived single cell clones from the SUM149PT ER^-^/PR^-^ inflammatory breast cancer cell line^21,22^ stratified by expression of CD44 and CD104 (ITGβ4)^14^. Six single-cell derived clonal populations were isolated, ranging from epithelial-like (E) to mesenchymal (M1 and M2), including three distinct intermediate states (EM1, EM2, and EM3) – hereafter referred to as “EMT clones” (Figure 1A, Supplemental Figure 1A). These clones, which stably retain their EMT states in culture, were ranked along the EMT spectrum relative to one another based on expression of canonical epithelial and mesenchymal markers such as *Vim* (Vimentin), *CDH1* (E-cadherin), *ZEB1*, and *SNAI1* (SNAIL) (Figure 1C&D), as well as by variable migratory and invasive characteristics *in vitro* (Figure 1B). In contrast to previous reports^23,24^, we observe an increase in migratory and invasive properties in the intermediate, but not mesenchymal clones, likely pointing to earlier discrepancies in discerning these two states from one another. Classic flow cytometry approaches employing 2-3 markers were insufficient to distinguish the intermediate states (Supplemental Figure 1D). By other measures, however, the intermediate clones exhibit differences in cell migration (Figure 1B), EMT marker expression (Figure 1C&D), cell morphology (Figure 1E) and expression of Vimentin and E-cadherin (Figure 1F). Interestingly, the three intermediate clones most closely resemble the characteristics of the parental line in migratory and invasive ability *in vitro* (Figure 1B), as well as their overall transcriptional profiles determined by unsupervised hierarchical clustering of Pearson’s correlation coefficients following RNA-sequencing (Figure 1G). Notably, there were no substantial genetic differences between clones E, EM1, EM2, EM3 and M1, with M2 exhibiting some *SNP* and *INDEL* variations (Supplemental Figure 1B&C), indicating that the phenotypic and functional differences between these intermediate clones are likely driven by non-genetic mechanisms^25^.

**Figure 1:**
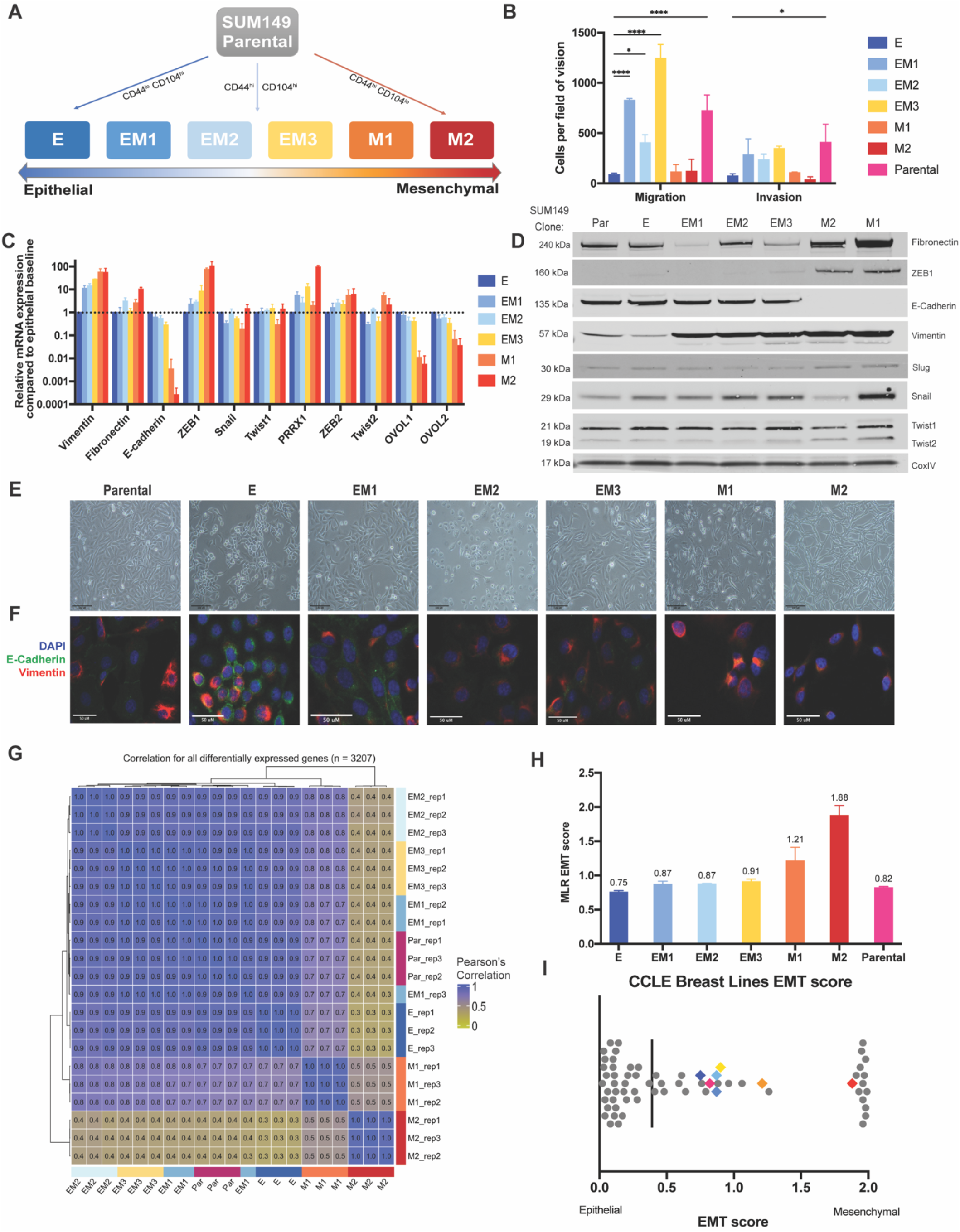
Heterogeneous cell line SUM149PT contains multiple distinct EMT states that can be isolated as single cell clones. A) A schematic of the Flow cytometry method used to isolate single cell clones that present as an epithelial (E), three distinct intermediate (EM1, EM2, and EM3), and two mesenchymal (M1 and M2) EMT states. B) *In vitro* assessment of clonal migratory and invasive characteristics as measured in a standard transwell assay (n=3, SD, p-value ***<0.001, *<0.05). Canonical EMT marker expression levels as determined by (C) Quantitative RT-PCR (SD, n=4) or (D) immunoblotting to rank SUM149 clones along the EMT spectrum. E) Bright field and F) immunofluorescent images of EMT clones *in vitro* stained with Vimentin and E-cadherin displaying cell morphology and marker expression & localization, respectively. G) Unsupervised hierarchical clustering of Pearson correlation coefficients of clone-specific gene expression profiles derived from RNA-sequencing. Correlation coefficients were determined using a total of 3207 genes, each of which was differentially expressed between epithelial clone, E, and at least one of the comparison clones. Yellow indicates low correlation whereas blue indicates higher correlations. H) EMT signature of EMT clones and parental line generated from the ordinal Multinomial Logistic Regression method of gene scoring and I) distribution of EMT score of the EMT clones among other breast cancer cell lines from the Cancer Cell Line Encyclopedia (CCLE).

Various methods have been developed to quantify the extent to which cells undergo an EMT and determine an absolute comparable EMT score^26–28^, which have been reviewed recently^29^. When applied to our model, all three methods - MLR^28^, KS^27^ and 76GS^26^ - predict that E through EM3, i.e. the epithelial and three intermediate clones, fall within the intermediate state, while M1 and M2 are mesenchymal (Figure 1H, Supplemental Figure 1E, F, & G). When plotted amongst the 59 breast cancer cell lines from the Cancer Cell Line Encyclopedia (CCLE), these clones fall along the intermediate and more mesenchymal end of the spectrum (Figure 1I, Supplemental Figure 1E&F, Supplemental Table 1). Notably, the parental line scored most closely to the epithelial (E) clone in most methods (Figure 1H, Supplemental Figure 1E&F). Our model, thus, highlights the presence of multiple distinct intermediate EMT states, enabling further investigation into how each state may individually contribute to tumor development and metastasis.

*In vivo* tumor initiation and growth further highlighted the individuality of these EMT clones. Upon orthotopic injection, the parental cell line was able to initiate and form tumors more rapidly than the other clones (Figure 2A). When analyzed with a bimodal linear mixed model fit^30^, the three intermediate clones were able to initiate tumors at the same rate as the parental line (Initial phase Holm adj. p-val > 0.05) (Figure 2A & 2C), but exhibited a lag in growth (Exponential phase Holm adj. p-val <0.003). The epithelial (E) and two mesenchymal (M1 & M2) clones both failed to initiate tumors as readily or, in the case of the mesenchymal clones, grow as rapidly as the parental and intermediate clones (Holm adj. p-value < 0.01). Increased tumor growth corresponded with decreased survival, with the parental line exhibiting poorest survival, and the mesenchymal clones the longest (Figure 2B). Limiting dilution analyses revealed a high tumor-initiating cell frequency in the parental and intermediate clones, with all mice forming tumors by eight weeks (Figure 2C). The two mesenchymal (M1& M2) possess the lowest tumor initiating cell frequencies (1/694,444 and 1/75,975, respectively) despite expressing high levels of CD44 expression, a marker frequently correlated with increased stemness and tumor initiating ability^11,12,31^ (Supplemental Figure 1D). All clones generated high grade, poorly differentiated, invasive ductal carcinomas of no special type (ductal), with clone M1 tumors notably exhibiting 50% squamous differentiation, and clone M2 showed abundant spindle cell morphology (Supplemental Figure 2A). Tumor growth and TIC frequencies indicate that, while the intermediate EMT clones may represent a population with high tumor-initiating potential, a heterogeneous population such as the parental line is able to enter an exponential growth phase more rapidly.

**Figure 2:**
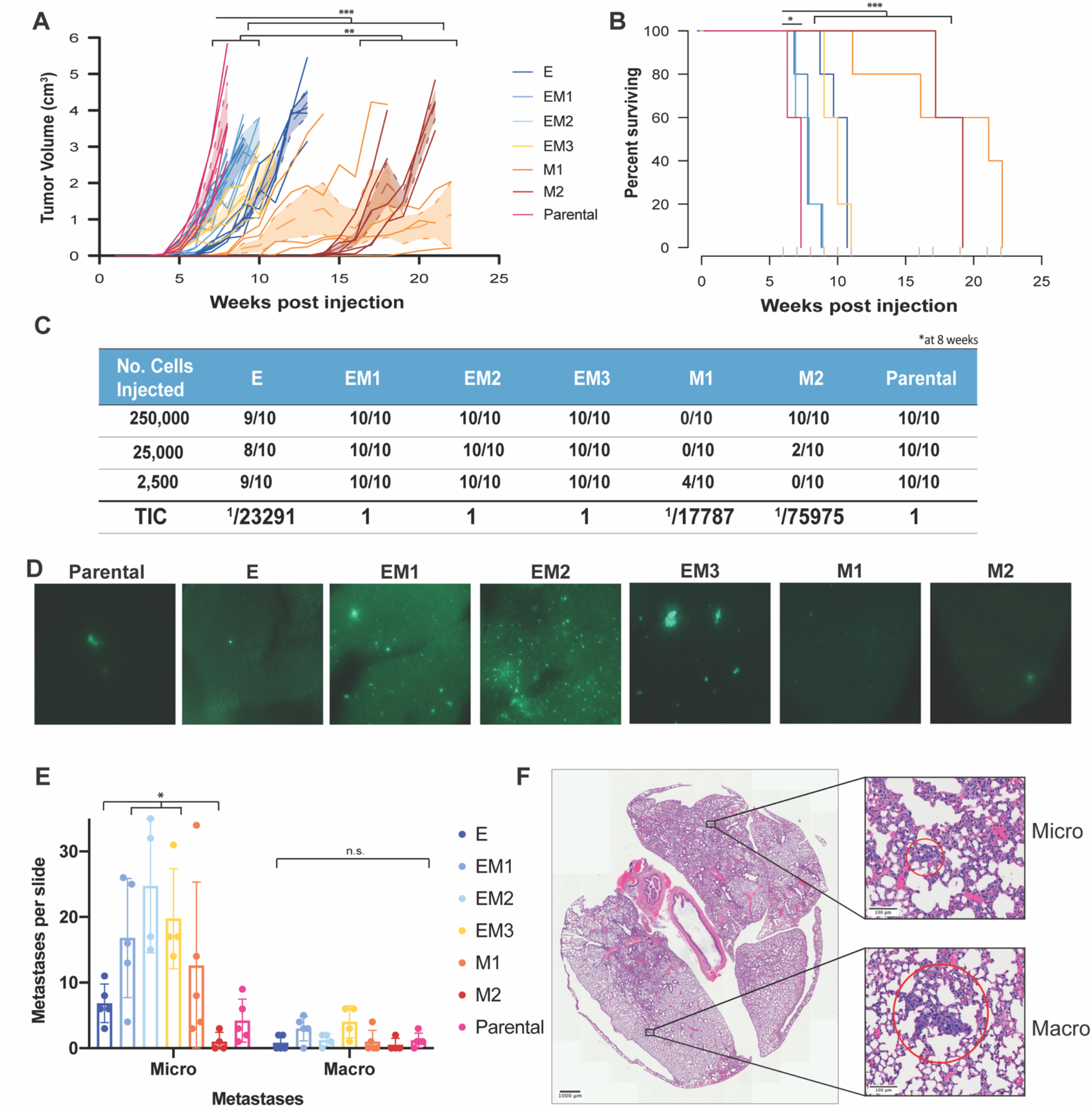
Differences in primary tumor growth and metastatic potential between EMT clones. A) Tumor growth curves measured weekly following orthotopic injection of clonal and parental cell lines at 2,500 cells exhibit exponential growth differences between EMT states (TumGrowth^30^ piecewise regression model breakpoint = 6wks, ***Holm adj. p-value <0.0001, **Holm adj. p-value< 0.01, n=10). B) Survival curve of EMT clones and parental line displaying differences in survival across the parental line and early intermediate EMT states. Cox regression analysis (***Holm adj. p-value <0.0001, *Holm adj. p-value< 0.015). C) Tumor initiating cell frequency calculated by limiting dilution assay with cells injected at 250k, 25k, and 2.5k cells per flank. TIC calculated at 8 weeks post injection. D) Lung sections collected at time of maximum tumor burden (2cm^3^), with GFP labeled metastatic cells, following orthotopic injections in (A). E) Lungs fixed and stained from (A) with H&E and enumeration of micro (<10 adjacent cells) and macro (10+ adjacent cells) metastatic regions (SD, n=5, micro p-val < 0.02 macro p-value = n.s.). F) Representative brightfield images of micro- and macro-metastases from the parental line.

To precisely delineate the propensity of the clones to form micro- and macro-metastasis, lungs from animals bearing orthotopic tumors were fixed and stained with H&E and counted for micro- (<10 adjacent cells) and macro- (>10 adjacent cells) metastases (Figure 2E). The three intermediate (EM1-3) clones seeded higher numbers of micro-metastatic lesions per lung, compared to the most epithelial (E) and most mesenchymal (M2) (p-value <0.05). Within this group, clones EM1 and EM3 seeded higher numbers of macro-metastases compared to EM2 (Figure 2E), although differences between the intermediate clones were not statistically significant. Interestingly, while exhibiting the highest rate of tumor growth and poorest survival, the parental line seeded fewer lung metastases than the intermediate clones, suggesting that other mechanisms could be contributing to mortality.

Given the lack of genetic differences between the clones, we hypothesized that the clonal variations in EMT state were driven by alterations to their transcriptional profiles. Using RNA-sequencing (RNA-Seq), alterations in the expression levels of various transcription factors were analyzed using the epithelial clone E as a benchmark. Principal Component Analysis (PCA) demonstrates clustering of the parental line, intermediate EM and E clones (Figure 3A, Supplemental Figure 3A), whereas the two mesenchymal clones (M1 & M2) cluster separately from each other and the main cluster, indicating that the EMT clones do not reside in a linear spectrum, but rather embark on multiple distinct trajectories. Unsupervised hierarchical clustering of the 500 most variable genes across all clones reveals distinct transcriptional programs separating the three intermediate clones from the epithelial (E) and mesenchymal (M1 & M2) ones (Supplemental Figure 3B). Within this intermediate cluster, EM2 and E are again distinct from the remaining intermediates and parental line (Supplemental Figure 3A). Differential expression analysis revealed 581 genes were significantly differentially expressed (adjusted p<0.05) expressed in all clones when compared to E, with 1881 genes shared between the two mesenchymal clones and 178 shared between the three intermediate clones (Figure 3B, Supplemental Figure 3C, Supplemental Table 2). Indeed, in comparison to the epithelial baseline clone E, more differentially expressed genes are exclusive to each clone than are shared between two or more clones (Figure 3B), further corroborating the unique EMT states represented by this model. To further explore the epigenetic landscape of these clones, we employed ATAC-seq^32^ to determine the differential chromatin accessibility across EMT clones in comparison to the epithelial clone E, which exhibits the most closed chromatin profile (Figure 3C). Notably, the chromatin landscape is similarly diverse between mesenchymal and intermediate clones (Supplemental Figure 3D). To identify TFs with a significant enrichment of motifs among peaks that were uniquely accessible in each clone (relative to E), presence of transcription factor binding motifs was scanned for using Motifmatchr^33^, and tested for enrichment against the background set of all identified peaks (hypergeometric test, p-adj<0.05) (Figure 3D). The three intermediate and early mesenchymal clones were highly enriched for motifs of the RUNX family transcription factors (Figure 3D, Supplemental Figure 4A). The TFAP2A/B/C TF family also exhibit enriched binding accessibility in these intermediate and early mesenchymal clones, albeit less significantly than the RUNX family. All three RUNX transcription factors as well as their co-factor CBFβ have been previously implicated in various cancers^34–36^ and in metastatic progression of triple negative breast cancer (TNBC)^37,38^. Enrichment of these RUNX TF motifs is unique to the intermediate (EM1, EM2, and EM3) and early mesenchymal (M1) clones, those that seeded the highest number of lung metastases (Figure 2E).

**Figure 3:**
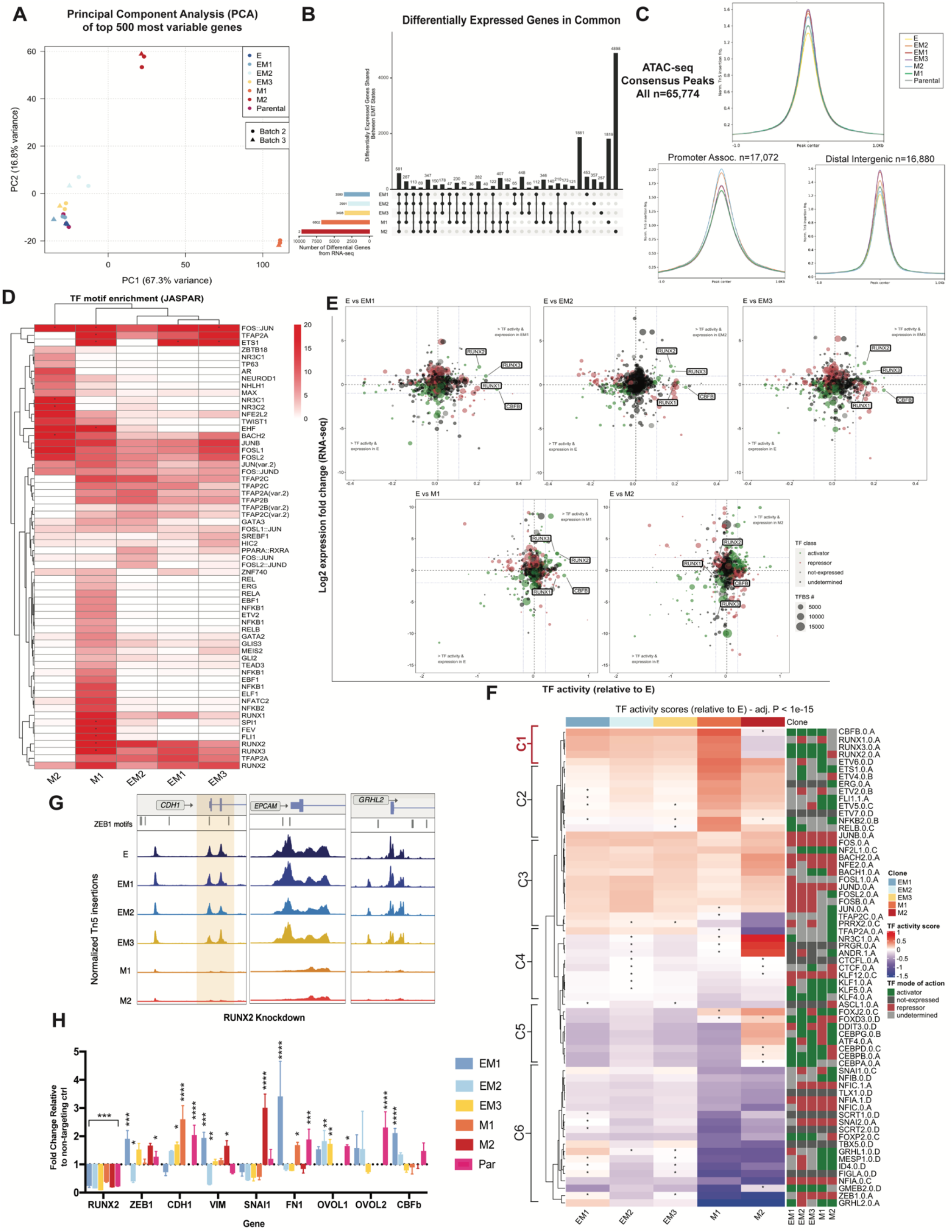
Identification of stabilizing transcription factors in the intermediate EMT state by multi-omics analysis. A) Principal Component Analysis (PCA) of the 500 most variable genes between all EMT clones and SUM149 Parental line from RNA-sequencing. B) C) ATAC-seq peak accessibility measured as counts per million (CPM) normalized Tn5 insertions surrounding consensus peaks, promoter-associated, and distal-intergenic peaks, respectively, for each EMT clone D) Unsupervised hierarchical clustering of transcription factor motif enrichment (-log10 adjusted p-value, hypergeometric test) among accessible chromatin peaks unique to each clone, relative to E. Presence of motifs obtained from the JASPAR database were identified using motifmatchr (P<0.05) and tested for enrichment against the background set of all peaks identified in the respective clone using the hypergeometric test. Asterisk indicates a -log10 p-value enrichment threshold greater than 20, scaled to fit. E) Advanced volcano plot of highly significant transcription factors, highlighting the RUNX family Transcription Factor activity, relative to clone E determined by diffTF from ATAC-seq along the X-axis (label cutoffs at 0.1, −0.1 TF activity), plotted against Log2 fold gene expression values of transcription factors on the y-axis (label cutoffs at 1, −1 log2 fold). Transcription Factor classification, determined by transcription factor expression, displayed in bubble color, and number of transcription factor binding sites used to determine TF activity plotted as bubble size. F) Unsupervised hierarchical clustering of TF activity scores (Z-score transformed) compared to clone E (adj. P-value <1E^-15^). Right hand side displays TF classification, determined by changes in TF expression (diffTF), as an activator, repressor, not expressed, or undetermined. G) Peak accessibility of Tn5 normalized, merged coverage of three canonical ZEB1 target genes, CDH1, EPCAM, and GRHL2 across all clones from ATAC-sequencing. ZEB1 TF motifs highlighted above signal tracks. H) mRNA levels of canonical EMT markers determined by qRT-PCR following inducible shRNA knockdown of RUNX2 at 72 hours post-induction in intermediate, mesenchymal, and parental clones. Fold change relative to a LacZ nontargeting control (dotted line at 1) (SD, n=4, * <0.01, **<0.005, *** < 0.001, ****< 0.0001, no star = n.s.)

To provide a more comprehensive picture of the epigenetic and regulatory landscape of EMT, we utilized a new multiomics approach, DiffTF^39^, to quantify TF activity and regulatory state by combining RNA-seq and ATAC-seq data for each clone. Similar to Motifmatchr, TF activity was inferred by computing the fold change in chromatin accessibility between each clone compared to E at each binding site of a given TF. Inferred TF activity was visualized against RNA-seq log2 fold-change for each transcription factor (Figure 3E, Supplemental Figure 3E), facilitating detection of TFs with increased expression in conjunction with increased TF activity (upper right quadrant). Analysis of the directional changes of TF expression from RNA-seq also indicated whether the TF in question acts as an activator (green) or repressor (red) (Figure 3E). The two mesenchymal clones revealed much larger shifts in both fold-change expression and TF activity, indicative of larger changes in their overall transcriptional and chromatin profiles relative to E. In the three intermediate and early mesenchymal clones, RUNX2 and RUNX3 are both amongst the most highly expressed and exhibit the highest TF activity of all significant transcription factors (Figure 3E, Supplemental Figure 3E). A list of all significant transcription factors and their relative TF activity and fold change expression are included in Supplemental Table 3.

Unsupervised clustering analysis presents a clearer picture of how these TFs act in a transcriptional network, highlighting those which exhibit significant changes in their TF activity in at least one clone (adj. p-value <1E^-15^) (Figure 3F). Cluster 1 is comprised of transcription factors that are upregulated and function as activators in the clones that exhibit increased metastatic potential i.e., EM1-3 and M1; the RUNX TF family and co-factor CBFβ. Cluster 2 includes factors that increase in activity with the progression of EMT e.g., ETS1 and related factors, NFkB, RELB, and FLI1. Cluster 3 identifies transcription factors that are activated and remain equally active after entrance into the EMT e.g., members and regulators of the AP1 complex. Multiple members of the KLF family of transcription factors as well as CTCF are found in cluster 4, which exhibit a consistent decrease in TF activity following entrance into an EMT. Cluster 5 highlights transcription factors that may be uniquely active in M2, while remaining inactive in the other EMT clones, including FOXJ2 and FOXD3, CEBP complex members, and ATF4. Finally, Cluster 6 delineates the transcription factors that have an overall and graded decrease in activity as the EMT progresses. This cluster contains multiple EMT TFs such as Snail, Slug, and ZEB1 as well as known MET promoting TFs such as GRHL1 and 2. Although these and other canonical EMT transcription factors, such as Twist and PRRX1 are expressed at a significantly high levels (p<0.05), non-significant or decreased TF activity indicates no change in chromatin accessibility compared to clone E in response to this increase in expression (Supplemental Figure 3E, Supplemental Table 3). Indeed, these TFs decrease in activity while their overall expression remains high (Supplemental Figure 3E) likely resulting from strong repression of chromatin accessibility at their epithelial target genes, potentially through HDAC recruitment^40,41^ or other mechanisms such as the ZEB1 target promoters *CDH1, EPCAM, GRHL2* (Figure 3G).

To validate our analyses, we tested the effects of knockdown of TFs in cluster 1, which is uniquely expressed and active in the intermediate and early mesenchymal clones. Indeed, knockdown of *RUNX2* led to a significant upregulation of *CDH1* in clones EM2, EM3 and M1, a reduction in *Snai1* and increase in *OVOL1* expression across all intermediate clones, indicating the acquisition of epithelial traits (EM1, EM2 & EM3; Figure 3H, Supplemental Figure 4B&C). In addition to validating our approach of identifying key regulators of the intermediate EMT states, RUNX2 presents itself as a viable candidate for potential disruption of this state.

To better understand the role that epithelial-mesenchymal (E-M) heterogeneity plays on tumor progression and metastasis, we utilized a multi-round, multiplexed tyramide signal amplification (TSA) approach^42^ to assess protein levels of EMT markers while segregating the epithelial component of tumors from stromal elements that could obfuscate EMT scoring. To capture the full spectrum of EMT states, we designed a panel of six EMT markers, containing three intermediate filament proteins (KRT8, KRT14 and VIM), two EMT transcription factors (ZEB1 and Snail), and an adherens junction protein that serves as a hallmark epithelial marker (E-cadherin). Tumors from each of the clones was stained and divided into 50 regions of interest (ROI) and processed with the InForm™analysis software (PerkinElmer) to generate composite images (Figure 4A). Image training algorithms were used to discern tumor and stromal composition, as well as cell segmentation (Supplemental Figure 5A). Normalized percentile distributions of the arbitrary fluorescent units (AFU) of each EMT marker, per cell, across all images provided an overview of the overall composition of these tumors (Figure 4B), indicating that no one unique marker signature defined any individual tumor. Overall, clone M2 tumors expressed less E-cadherin and more ZEB1, while clone E tumors contain less Vimentin, indicative of their initial EMT states in-vitro. To determine the extent of E-M heterogeneity within the stained images, a scoring metric was developed using the SUM149PT clones as a training set. Tumor images were scored based on a rubric of low (one major cell trait with up to one minor trait), mid (two major cell traits with up to three minor traits), and high (three or more major cell traits present with two or more minor traits) (Figure 4C). Scoring based on these criteria revealed that the intermediate EMT clones form tumors that contain more regions of higher heterogeneity than the parental cell line (Figure 4C). Thus, increasing levels of heterogeneity may not linearly correlate with tumor growth or metastasis, but rather, there exists an optimal ratio of cell traits within the tumor that determines its growth and metastatic potential. Indeed, the requirement for this optimal ratio may explain the growth lag observed in the intermediate clones when compared to the parental line, which likely results from constraints in the generation of a requisite level of heterogeneity from a homogenous cell culture (Figure 2A).

**Figure 4:**
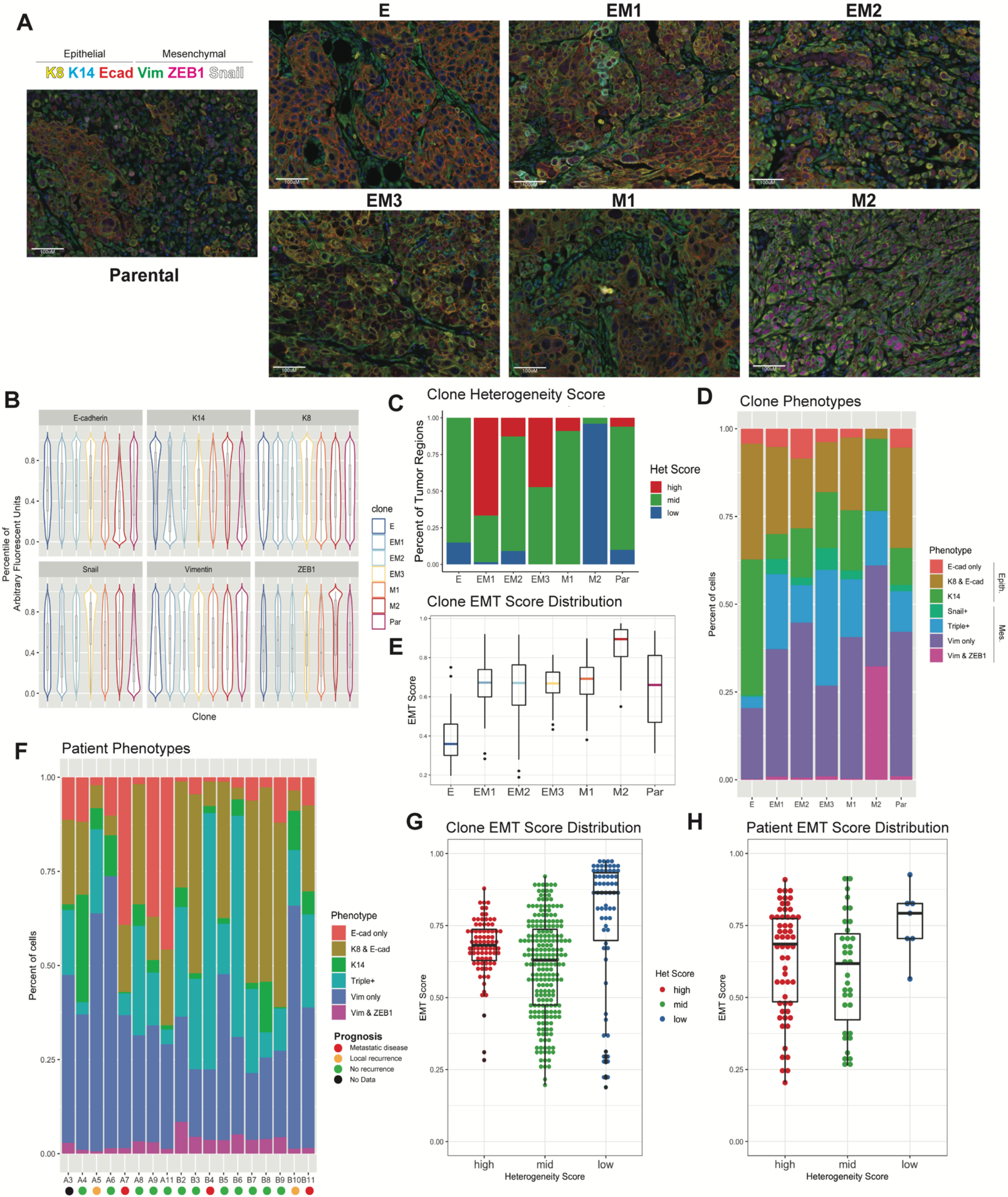
Multiplexed staining of tumors reveals shared phenotypes also found in patient tumors. A) EMT clone-derived tumors resected at 1.5cm^3^ and stained with six markers using TSA (n = 50 images) B) Percentile normalized distributions of Arbitrary Fluorescent Units (AFU) for each marker across all tumors. C) Empirically determined heterogeneity scores of EMT clone-derived tumors. Rubric: Low (one major cell trait with up to one minor trait), Mid (two major cell traits with up to three minor trait), and High (three or more major cell traits present with two or more minor traits). D) Boxplot of EMT phenotypes determined from InForm Analysis software displaying the composition of each clonally derived tumor (n=~50 images per tumor). E) EMT score distribution in clonally derived tumors generated from weighted multivariable logistic regression of the phenotypes present in each tumor. F) Phenotypic composition of 20 patient tumors (n=3-7 punches per tumor) stained with the TSA marker panel. Colored dots correspond to patient prognosis, either No Recurrence, Local metastasis, metastatic disease, or no data G) Dotplot of correlations between heterogeneity score and EMT score in SUM149PT clone tumors and H) matching pattern in patient tumors.

In a parallel method to assess tumor diversity and phenotypic composition, the InForm analysis software was trained on a subset of tumor images to recognize seven distinct cell phenotypes encompassing the majority cell states present in all tumor images (Supplemental Figure 5A). These phenotypes, spanning from most epithelial to most mesenchymal are: E-cadherin only, KRT8 & E-cadherin, KRT14 only, Triple positive (KRT8+E-cadherin+Vimentin), Snail only, Vimentin only, and Vimentin+ZEB1. Phenotypes were validated manually and by fluorescent marker distribution (Supplemental Figure 5B). Similar to EMT marker distribution, there were no phenotypes unique to any individual state, indicating that optimal tumor progression may be determined by ratios of cell states rather than by presence or absence of a particular cell type. On either end of the spectrum, clone E and clone M2 derived tumors are made up of over 75% epithelial (E-cad only, KRT8+E-cad, or KRT14 only), or mesenchymal (Snail only, Vimentin only, or Vimentin+ZEB1) phenotypes, respectively, whereas tumors derived from the intermediate clone and the parental line contain equal distribution of epithelial and mesenchymal phenotypes (Figure 4D). These data further support the notion that the ratio of EMT states (phenotypes), rather than the presence of any particular state, is more reflective of tumor growth and metastatic potential.

In order to develop a scoring algorithm to assess E-M heterogeneity, three feature extraction methods were tested using the segmented cell data files (Supplemental Figure 5A). These three approaches sought to determine the best method to assess heterogeneity: (1) an entropy-based approach^43,44^ using mean marker expression of cells per image, (2) a nearest neighbor analysis approach^45^ using cell phenotypes as determined above (Figure 4D), and (3) a hybrid approach combining approaches 1 and 2. All three approaches were developed and evaluated using a randomized subset of clone tumor images and their corresponding heterogeneity scores according to the rubric outlined in Fig 4C as ground truths. Logistic regression performance on unseen test data indicated 78%accuracy for entropy-based features of mean marker cell expressions. Fivefold cross-validation determined that the entropy-based approach 1 (F1-score= 0.78, Wilcoxon ranking test p-value=0.004) proved to be the most accurate at correctly assessing tumor heterogeneity (Supplemental Figure 5C). This shows that E and M2 clones consisted largely of areas of low or mid-level heterogeneity, whereas all intermediate clones were comprised of regions of high heterogeneity (Figure 4C, Supplemental Figure 5C).

In addition to scoring E-M heterogeneity, we sought to assign EMT scores to tumors based on the ratios of epithelial and mesenchymal traits they exhibit. EMT scores were generated by calculating the composition of cell phenotypes per image (50 images per clone) using the linearly weighted average of cell ratios expressing each phenotype from epithelial to mesenchymal. EMT scores range from 0 to 1, with zero being comprised of all epithelial phenotypes and one comprised of all mesenchymal phenotypes. Rather than attaining an equilibrated EMT state, clonal tumors held true to the EMT state of their starting populations, with clone E tumors being predominantly epithelial (mean=0.35), intermediate (EM1, EM2, EM3) and parental tumors maintaining an intermediate EMT score (mean=0.65-0.7), and the most mesenchymal (M2) scoring >0.8 (mean=0.86) (Figure 4E). Clone M1 tumors also scored as intermediate despite starting as a quasi-mesenchymal, further validating this clone as a late intermediate (Figure 4E). Interestingly, the variation in EMT scores between images in the intermediate clones (EM1, EM2, EM3, & M1) and parental line was highest of all of the groups, indicating higher intra-tumoral heterogeneity (Figure 4E). In exploring the connection between heterogeneity and EMT scores, we uncovered that low heterogeneity tumor regions correlated with higher EMT scores, as can be seen in M2 tumors (Figure 4A & 4G, Supplemental Figure 5D). Medium and high heterogeneity regions were characterized by intermediate EMT scores despite encompassing a more diverse array of possible EMT states (Figure 4G).

To establish whether the EMT phenotypes observed in our model are representative of those observed in human breast cancer specimens, we carried out TSA immunostaining of 20 TNBC patient tumors (Supplemental Figure 5E & F) selected for TNBC status and poorly differentiated histopathology. Six out of seven phenotypes identified from the SUM149PT clone tumors were reproducibly represented across all patient tumors, with the exception of Snail only (Figure 4F, Supplemental Figure 5E). Ratios of epithelial and mesenchymal phenotypes follow similar trends to the SUM149PT clones, skewing towards more mesenchymal cells (Supplemental Figure 5G). These data serve to validate the SUM149PT clones as a model to study various states along the EMT spectrum. Heterogeneity score and EMT score algorithms were applied to these patient tumors (Supplemental Figure 5G, H, & I) and, as observed in the EMT model clones, relative intra-tumoral heterogeneity and EMT scores were not always correlated e.g., patient tumor A5 and A8 exhibit similar ratios of low, mid, and high intra-tumoral heterogeneity, however, their EMT scores are markedly different (Supplemental Figure 5G&H), which suggests that these two metrics may determine patient outcomes in different ways. This uncoupling of heterogeneity and EMT scores was also observed in the SUM149PT parental tumors compared to those from clone E which exhibit similar ratios of low and mid-level heterogeneity but are scored differently on the EMT scale. The observed uncoupling of heterogeneity from EMT state raises the notion that cells which exhibit high plasticity leading to the formation of heterogeneous tumors do not reside in a highly mesenchymal state, but rather, fall in the mid-range of EMT scores. Thus, TFs identified in cluster 1 such as RUNX2 which are associated with the intermediate state likely regulate cell plasticity, while not directly contributing to a more mesenchymal state.

Our study encompasses a comprehensive analysis of the spectrum of EMT states represented within a single cell line to interrogate the nuances of each state, their respective contributions to tumor initiation and metastasis, and their epigenetic regulatory networks. We uncover the presence of multiple unique EMT states within the intermediate EMT category, as well as two distinct mesenchymal-like states, suggesting multiple, nonlinear trajectories for EMT. The intermediate EMT state is maintained by a unique set of transcription factors, including the RUNX family, which are necessary for its stability. We also uncover that the intermediate EMT state exhibits higher levels of cellular plasticity, manifesting in tumors as high levels of E-M heterogeneity. Moreover, this plasticity-induced heterogeneity plays a key role in the metastatic propensity and tumor-initiating potential of these clones. Finally, we develop an approach to quantifying E-M heterogeneity and “EMTness” within human tumors, the former showing promise in its ability to inform disease prognosis.

## Materials and Methods

### Cell culture

The human derived SUM149PT cell line was obtained from the Weinberg lab, who in turn acquired it from Dr. Stephen P. Ethier (Michigan). All derivative cell lines were maintained in F12 medium (Gibco #11765-054) supplemented with 10% fetal bovine serum (FBS) (Gibco #10438-026) insulin (1 mg/ml; Gibco #12585-014), and hydrocortisone (1 mg/ml; Sigma # H4001) and 5% Penicillin-Streptomycin (Corning #10-002-cl). All cell lines were incubated at 37°C with 5% CO_2_-air atmosphere with constant humidity. Cells were passaged with 0.25% Trypsin (Corning 25-053-Cl); passage number was kept on all cell lines, and cultures were discarded past a total of twenty passages to maintain their respective EMT phenotypes. 293T cell line was maintained in DMEM + 10% fetal bovine serum (Gibco #10438-026) + 5% Penicillin-Streptomycin (Corning #10-002-cl).

### Lentiviral Vectors

Lentivirus was made with 293T cells plated at 60% confluency in 10cm tissue culture treated plates and transfected using X-tremeGENE HP DNA Transfection Reagent (Sigma # 6365779001) with lentiviral vectors, psPAX2 (1.5ug, Addgene #12260), pCMV-VSV-G (1.5ug, Addgene #8454), and pcDNA3-eGFP (0.5ug, Addgene #13031) plus the lentiviral vector of interest (3ug). Supernatant collected at 48 and 72 hours post-transfection, concentrated using Lenti-X concentrator (TakaraBio #631232), and titer determined with Lenti-X GoStix (TakaraBio #631243).

### Cell line Generation

Parental cell line SUM149PT was maintained in standard media. To generate single cell clones, fluorescence assisted cell sorting (FACS) was performed on SUM149PT with the FACSAria III Cell Sorter. Cells were stained with CD44-PeCy7 1:100 (Biolegend 103030), CD104-APC 1:200 (Invitrogen #50-1049-82), or Epcam-BV510 1:100 (BioLegend #324235) for 30 minutes on ice before addition of DAPI 1:1000 (SigmaAldrich 10236276001, 10mM stock). Gating and compensation were done on single stained controls, and cells were sorted into collection tubes and immediately plated at a dilution of 0.5 cells per well into a 96-well plate. Single cell clones were then expanded and assessed for EMT characteristics.

#### ZsGreen expressing cells

All SUM149PT clones and parental line were infected with high titer pHIV-Luc-ZsGreen (Addgene #39196) virus so as to generate ZsGreen and Luciferase expressing tumor cells for metastasis tracking in mouse. 6×10^5^ cells were infected in 6-well plates with 125uL of high titer virus in standard media with 5ug/mL Polybrene (Sigma-Aldrich). Media was changed after 24 hours, and cells allowed to expand for 48 hours before sorting for ZsGreen positive population on the FACSAria III Cell Sorter, as above.

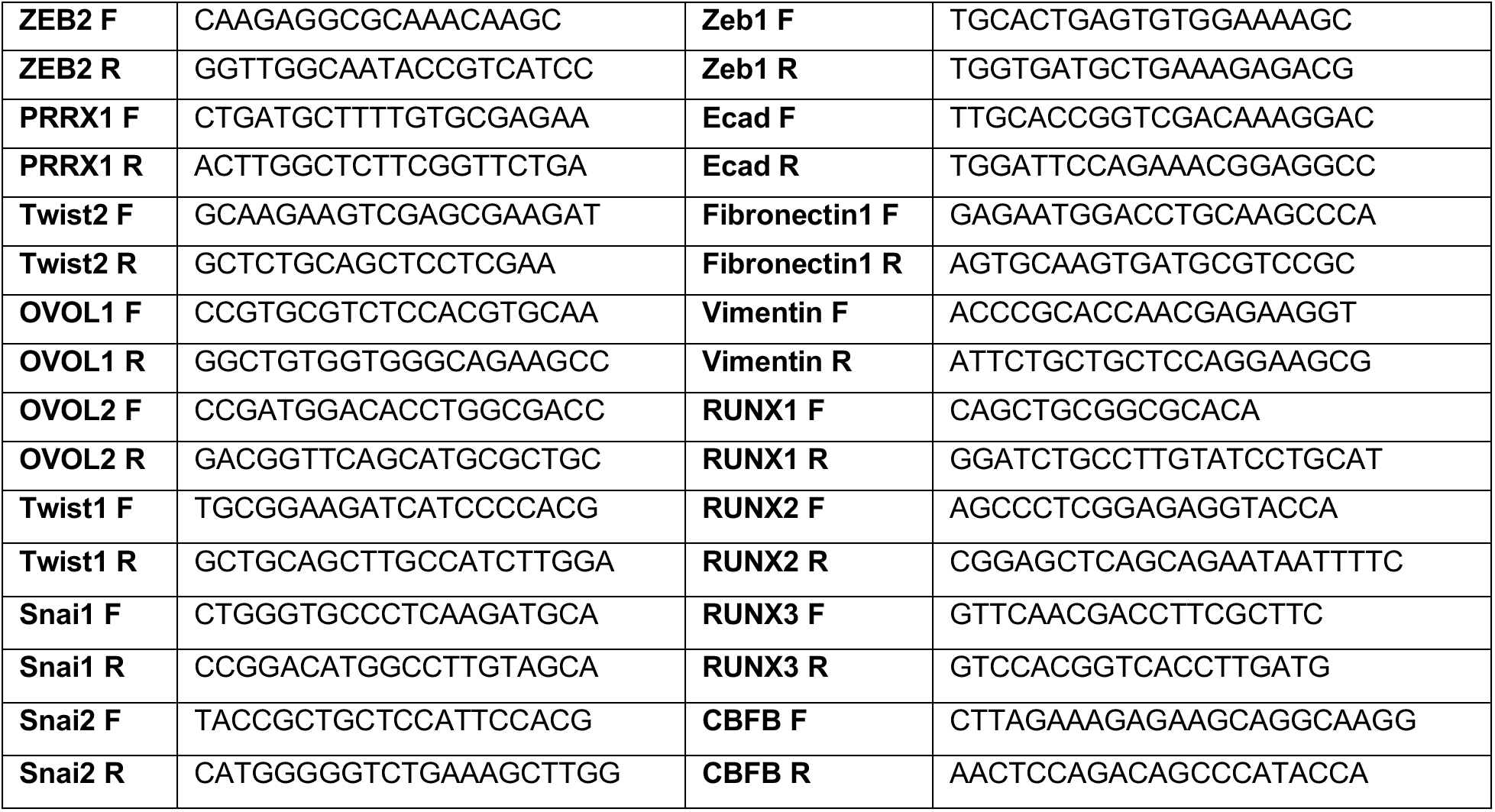

### Flow Cytometry

Flow experiments were performed in the same manner as above, on a 10-color Gallios FACS cytometer (Beckman Coulter). Compensation, file analysis, and plot generation was conducted using FlowJo (BD).

### RT-qPCR

RNA was harvested from 6-well plates of cells at confluency, extracted using the Qiagen RNeasy plus kit (Qiagen 74034) and quantified using a NanoDrop (Thermo Fisher Scientific - ND-2000-US-CAN). Reverse transcriptase PCR (AppliedBiosystems #4368814) was performed to generate cDNA and Power SYBR green PCR mastermix (Applied Biosystems) was used for qPCR.

### Western Blot

For western blot, lysates were collected on-plate with 1x RIPA buffer (EMD Millipore #20-188) with protease and phosphatase inhibitors (ThermoScientific #1861280). Lysates were sonicated and cleared before quantification with a Bradford Protein assay (BioRad) and loaded at 50ug per lane and run on a NuPage Bis-Tris gel (ThermoFisher) and transferred to nitrocellulose membrane with the iBlot semi-dry transfer system (ThermoFisher) and blocked in 5% milk in TBST before staining with Fibronectin (BD #610078, 1:10,000), ZEB1 (LSBio ## 1:2000), E-cadherin (BD #610182, 1:1000), Vimentin (Cell Signaling Technology; CST #5741, 1:2000), RUNX1 (CST #4336, 1:2000), RUNX2 (CST #12556, 1:2000), RUNX3 (CST #9647, 1:2000), CBFb (Abcam ab33516, 1:2000), Snail (CST #3879, 1:1000), Twist1/2 (abcam ab50887, 1:50), and CoxIV (CST #11967, 1:2000) overnight in 5% milk in TBST. LiCor secondary antibodies, IRDye goat anti-rabbit and goat anti-mouse, 800CW (LiCor #925-32219) and 680RD (LiCor #925-68076), applied at 1:10,000 for 1 hour RT in 5% milk in TBST before imaging on the LiCor Oddessey CLx Digital Imager.

### Transwell Assay

Transwell assays were conducted using Costar Transwell plates (#3422 8.0 μm) in triplicate. For migration assays, 2.5E^5^ cells were added to the top of each well in 10% complete medium, with 100% complete medium beneath the transwell. Cells were incubated for 16-18 hours at 37°C. Media was aspirated and cells were permeabilized with 100% methanol for 5 minutes RT followed by staining with crystal violet (0.5% crystal violet in 20% methanol). Transwells were imaged on a Nikon Eclipse TS100 and migrated cells were counted using ImageJ. Invasion assays were conducted as above on plates that were coated with 100uL of 5% Matrigel to the top of each transwell and allowed to set at 37°C for two hours before seeding.

### shRNA knockdown

RUNX2 knockdown was achieved by cloning RUNX2 hairpins (below) into Tet-pLKO-puro (addgene #21915) via AgeI and EcoRI. Constructs were confirmed by Sanger sequencing and used as vectors in lentivirus production as stated above. Cells were infected with 1/8^th^ lentiviral product from a 10cm plate in 6-well plates. Media was changed after 24 hours and puromycin selection (2ug/mL) was applied at 48 hours until stable cell lines generated.

Cells were plates at 2×10^5^ cells per well in 6-well plates and treated with doxycycline (2ug/mL) for 72 hours before on-plate lysis for RNA of protein collection.

### Cell Imaging

#### Bright Field

Images were taken on a Nikon Eclipse TS100 under 20x magnification to determine cell morphology

#### ImmunoFluorescence

Cells were grown to 60% confluency in chamber slides (Falcon 354118) with standard media. Cells were fixed and permeabilized prior to staining with primary antibody (Vimentin CST # 1:100 and E-cadherin BD number 1:100) overnight followed by secondary (anti-rabbit ThermoFisher #31466 1:10000, anti-mouse ThermoFisher #31431 1:10000) for one hour. Slides were washed and mounted using ProLong Diamond (Invitrogen P36961) before imaging on Zeiss LSM 800 with Airyscan (63X)

### *In Vivo* Studies

Cell lines were resuspended in 30% Matrigel (VWR 47743-706) and injected in limiting dilutions (250k, 25k, and 2.5k cells per flank) orthotopically into the inguinal mammary fat pat (no. 4) of NOD-scid IL2Rγ^null^ (Stock No: 005557; Jackson Laboratory). Tumor growth was monitored weekly and tumor volume was measured along 3 axes with calipers (VWR 62379-531). Tumors and lungs were harvested at time of tumor burden (total tumor volume of 2cm^3^) and fixed overnight with 10% neutral buffered formalin. Tumor growth curves and survival statistically analyzed using TumGrowth^30^. Tumor initiating potential calculated with Extreme Limiting Dilution Analysis (ELDA)^46^.

### Tumor and Lung staining

Tumors and lungs were extracted at time of tumor burden (total tumor volume of 2cm^3^) and fixed overnight with 10% neutral buffered formalin. All samples were then processed and stained for Hematoxylin & Eosin (H&E) by the Dartmouth Hitchcock Pathology Shared Resource. Lungs bearing ZsGreen positive EMT clones were counted by eye on Nikon Eclipse TS100 and select images taken at 4x magnification.

### Metastasis counting

H&E stained lung slides were scanned on a PerkinElmer Vectra3 slide scanner at 10x and counted by eye for micro (> 10 adjacent cells) and macro (10+ adjacent cells) metastatic tumors.

### Whole Exome Sequecing

Whole Exome sequencing and subsequent SNP and INDEL alignment and discovery was performed by BGISeq on all EMT clones. They obtained 9,242.46 Mb raw bases. After removing low-quality reads we obtained averagely 91,977,846 clean reads (9,197.79 Mb). The clean reads of each sample had high Q20 and Q30, which showed high sequencing quality. The average GC content was 50.63%. Reads were aligned with Burrows-Wheeler Aligner (BWA). The HaploTypeCaller of GATK(v3.6) was used to call and identified 190,503 SNPs and 33,385 InDels between all samples. SNPs and InDels for each clone were tested against the consensus set for clone E to determine possible genetic mutations with a Fishers Exact Test and plotted as Odds Ratios.

### RNA-seq data processing

RNA was collected using Qiagen RNeasy plus kit (Qiagen 74034) and quantified using a NanoDrop (Thermo Fisher Scientific - ND-2000-US-CAN).

Quality of raw single-end RNA-seq data was confirmed using FastQC^47^ (v0.11.8) before read trimming of polyA sequences and low quality bases using Cutadapt^48^ (v2.4). Reads were aligned to human genome hg38 using STAR^49^ (v 2.7.2b) with parameters “--outSAMattributes NH HI AS NM MD --outFilterMultimapNmax 10 -- outFilterMismatchNmax 999 --outFilterMismatchNoverReadLmax 0.04 --alignIntronMin 20 --alignIntronMax 1000000 --alignMatesGapMax 1000000 --alignSJoverhangMin 8 --alignSJDBoverhangMin 1”. Quality of alignments was assessed using CollectRNASeqMetrics (Picard Tools^50^) and duplicate reads were identified (but retained) with MarkDuplicates (Picard Tools^50^). Gene-level abundance estimates were generated using RSEM^51^ (v1.3.2) using the rsem-calculate-expression command with the parameters “--strandedness reverse --fragment-length-mean 313 --fragment-length-sd 91”.

### Downstream RNA-seq data analysis

Gene-level abundance estimates generated with RSEM were imported into R and analyzed using R-package DESeq2^52^. Tp perform exploratory analysis of global transcriptional profiles, abundance were transformed using the regularized logarithm approach^53^ implemented in R-package DESeq2, and the top 500 most variable genes across all clones were supplied to the prcomp() command in R to perform principal components analysis (PCA). The 500 most variable genes were also used to perform Unsupervised hierarchical clustering with the R-package ComplexHeatmap. Differential expression analysis was performed on the raw gene-level abundance estimates assuming a negative binomial distribution, with clone E used as the reference group in all comparisons. Gene-wise dispersion estimates were reviewed in all analyses to confirm the selected model was an appropriate fit for the data. Genes with a Benjamini-Hochberg adjusted P-value < 0.05 (Wald-test) were considered statistically significant.

### ATAC-seq data processing

Tagmented DNA and library prep for ATAC-sequencing was performed according to the protocol detailed in Buenrostro et. al. 2015^32^

Prior to analysis, quality of raw DNA sequences (in FASTQ format) was confirmed using FastQC^47^ (v0.11.8). ATAC-seq data was then processed using the publicly available ENCODE ATAC-seq pipeline (https://www.encodeproject.org/pipelines/ENCPL792NWO/), and relevant commands and options used are described in detail below. Illumina adapter and transposase sequences were trimmed using Cutadapt^48^ (v1.9.1) with parameters “--minimum-length 5 -e 0.1”. Trimmed reads were then aligned to human genome hg38 using Bowtie2^54^ (v2.2.6) in “--local” mode with parameters “-X 2000 -k 2”. Duplicate reads were identified using MarkDuplicates (Picard Tools^50^) and filtered from final alignments, in addition to unmapped reads and reads aligning to mitochondrial DNA, retaining only alignments formed by properly paired reads. For multi-mapping reads, one paired-end alignment was randomly selected as the primary alignment while the remaining alignments were discarded. Alignments (in BAM format) were converted to tagAlign files and shifted +4 bp and −5 bp on the + and – strands, respectively, to account for insertion of adapter sequences by Tn5 transposase. Peaks were called for each replicate using the MACS2^55^ (v2.1.1) callpeak command with parameters “--shift −75 --extsize 150 --nomodel --keep-dup all --call-summits -p 1.0E-10” and filtered against the ENCODE hg38 blacklist. The Irreproducible Discovery Rate (IDR) method was used to identify a set of reproducible peaks across biological replicates using an IDR threshold of 0.05. For visualization purposes, position-shifted (to account for Tn5 insertion) BAM files for biological replicates were merged MergeSamFiles (Picard Tools^50^) and used to generate counts per million (CPM)-normalized signal-tracks (in BigWig format) via the deepTools^56^ (v3.4.3) bamCoverage command with parameters “--binSize 5 --normalizeUsing CPM --effectiveGenomeSize 2913022398 --ignoreForNormalization chrX”. Heatmaps of normalized Tn5 insertions in peak regions or specific regions were generated using deepTools commands computeMatrix and plotHeatmap.

Several metrics were used to confirm ATAC-seq data quality. To confirm sequencing libraries were of sufficient complexity, three specific quality control metrics were evaluated: non-redundant fraction (NRF, number of uniquely mapping reads / total read number), PCR bottlenecking coefficient 1 (PBC1, number of genomic positions with at least 1 read mapped / number of distinct genomics position to which a read maps uniquely), and PCR bottlenecking coefficient 2 (PBC2, number of locations where one read maps uniquely / number of genomic regions where two reads map uniquely). Fragment length distributions were generated and reviewed in R using the “Rsamtools” package. The fraction of reads in nucleosome-free regions (NFRs) was calculated to confirm a sufficient fraction of reads were located in NFRs. Fraction of reads in peak regions (FRiP score) was calculated to assess the quality of the final IDR peak set. Enrichment of accessibility over transcriptional start sites (TSSs), calculated as the maximum number of normalized Tn5 insertions across a +/- 2kb region flanking hg38 TSS regions, was used to was to confirm data quality in a peak agnostic fashion.

### Downstream ATAC-seq data analysis

Basic peak annotation was performed using the annotatePeak() function from the ChIPseeker^57^ package, using a range of +/- 3kb to define promoter-associated regions. R-package TxDb.Hsapiens.UCSC.hg38.knownGene was used to define gene models and coordinates of genomic features. To create a set of consensus peaks, the IDR peak sets for each sample groups were merged using the GenomicRanges R-package. Tn5 insertions occurring in each peak of the consensus peak set were counted from position-shifted BAM files using the featureCounts() function (from the Rsubread package) with options “isPairedEnd=TRUE, countMultiMappingReads=FALSE”. To perform exploratory analyses of global chromatin accessibility profiles, raw counts were transformed using the regularized logarithm approach^53^ implemented in R-package DESeq2. Principal components analysis (PCA) was performed on the 3000 most variable consensus peak regions using the prcomp() function in R. The most variable peaks were defined as those with the greatest standard deviation across all samples. Unsupervised hierarchical clustering was performed with the ComplexHeatmap R-package, also using the 3000 most variable consensus peaks. Differential accessibility analysis of the consensus peak was also performed using the DESeq2 R-package, modelling raw counts using a negative binomial distribution, with clone E used as the reference group in all comparisons. Peaks with a Benjamini-Hochberg corrected P-value < 0.05 (Wald-test) were considered statistically significant.

### Differential TF activity analyses

Differential TF activity between single-cell derived clones, as well as TF mode of action (i.e. activator, repressor) was estimated using diffTF^39^. Briefly, when used with ATAC-seq data, diffTF computes the fold change in chromatin accessibility between two conditions at each binding site of a given TF, and the distribution of fold changes is compared to a set of background fold-change values to assess statistical significance of differences in TF activity between the conditions. DiffTF was used in conjunction with matched RNA-seq (classification mode) to classify each TF into one of the following modes of action (activator, repressor, not-expressed, undetermined) through correlation of TF expression levels with target site accessibility. DiffTF was used with options “pairedEnd” and “RNASeqIntegration” set to ‘true”, with all remaining options using default settings. *In-silico* predicted binding sites based on the HOCOMOCO v11 database^58^ and PWMScan^59^ for hg38 across 768 human TFs was used to define the atlas of TFBS for diffTF analyses. To concentrate on the TFs with the most confidently estimated TF activity scores (weighted_meanDifference), we restricted our downstream analysis to TFs that achieved an adjusted P-value of 1E-15. To identify modules of cooperatively regulated TFs, unsupervised hierarchical clustering was performed on the diffTF activity scores using R-package ComplexHeatmap.

### Enrichment of TFBS in clone-specific peak sets

To identify potential TFs responsible for mediating clone-specific phenotypes, we tested the differentially accessible peaks between clone E and each clone for over-representation of TF binding site motifs. We first restricted each peak set to regions that demonstrated statistically significant increases in chromatin accessibility compared to clone E (see description of differential accessibility analyses above) and scanned these peaks for TF motif occurrences using R-package motifmatchr^33^. Position frequency matrices for human TF motifs used as input to motifmatchr were downloaded from the JASPAR database^60^ using R-packages JASPAR2018 and TFBSTools^61^. Over-represented TF motifs in each peak set were identified through hypergeometric testing using the R function phyper(), with all peaks identified in that clone used as the background set. TF motifs with a Bonferroni corrected hypergeometric *P*-value <0.05 were deemed as overrepresented. To identify potential groups of coordinately regulated TFs across the respective clones, -log10-transformed *P*-values from hypergeometric testing were subjected to hierarchical clustering and visualized using R-package pheatmap. To prevent extreme motif enrichments from dominating the heatmap scale, -log10-transformed *P*-values were capped at a maximum value of 20 (highlighted with an asterisk).

### Multiplexed TSA staining

Tumors were selected from each EMT clone at approximately 1cm^3^ and stained with (in order) Snail (CST #3895, 1:400), KRT8 (Invitrogen PA5-29607, 1:300), KRT14 (Invitrogen MA5-11599, 1:1000), Vimentin (CST #5741, 1:500), E-cadherin (BD #610182, 1:500), and ZEB1 (Invitrogen PA5-82982, 1:1000). Antibody optimization and multiplexed staining was done according to PerkinElmer OPAL Assay Development Guide (August 2017) and previous literature^42,62^. Briefly, slides were baked to remove paraffin wax, and then sequentially washed with Xylene, rehydrated with decreasing concentrations of ethanol, and finally ddH20 before blocking. Then slides were incubated with primary antibody then secondary antibodies for 30 minutes at RT. Following washes, the selected OPAL fluorophore was applied to slides for precisely 6 minutes at RT in the dark, washed off, and slides microwaved at 20% power for 15 minutes to affix OPAL to target regions and remove primary and second antibody. Slides were blocked again and staining process repeated for each marker, and finally spectral DAPI (PerkinElmer, 2 drops per mL) before mounting on coverslips with ProLong Diamond (Invitrogen P36961).

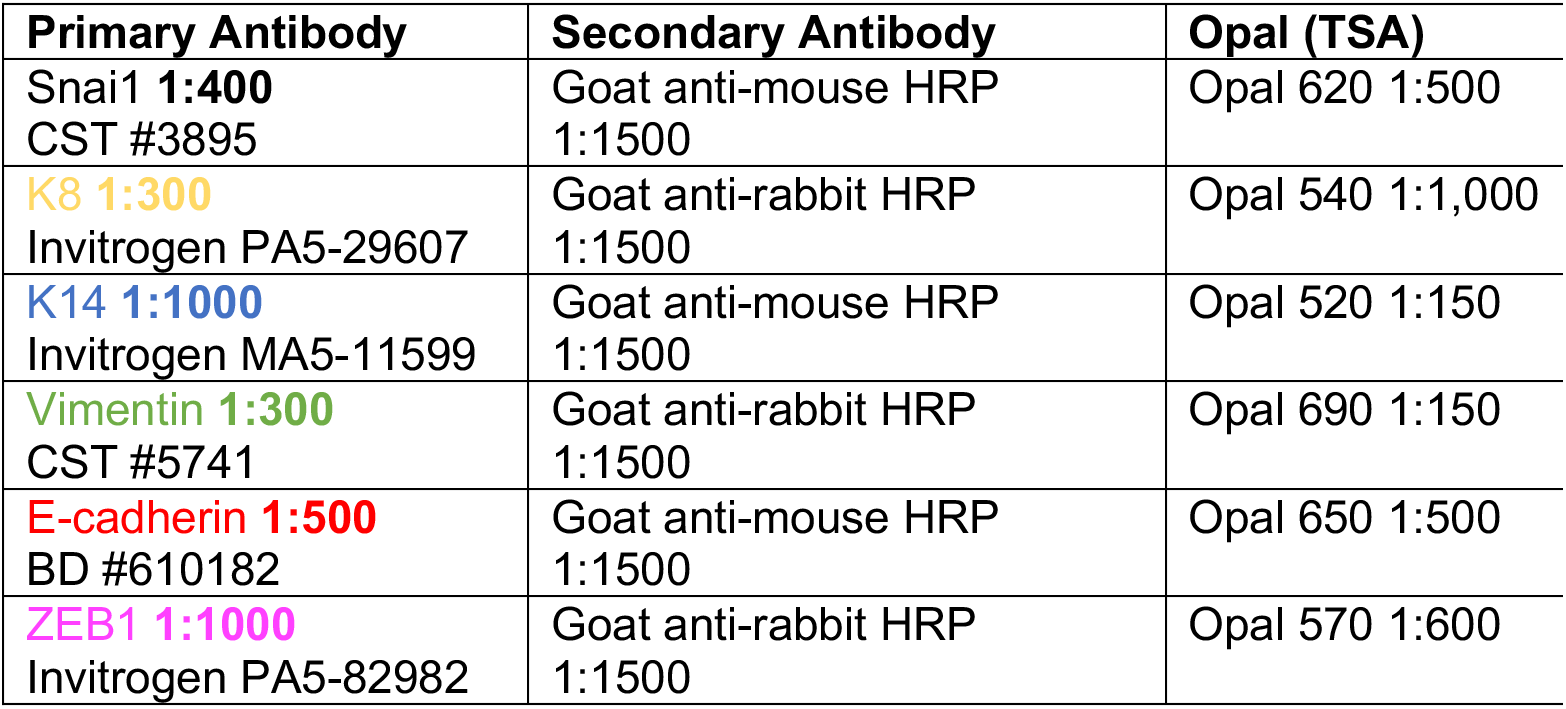

| **Primary Antibody** | **Secondary Antibody** | **Opal (TSA)** |
| --- | --- | --- |
| Snai1 **1:400**<br>CST #3895 | Goat anti-mouse HRP<br>1:1500 | Opal 620 1:500 |
| **K8 1:300**<br>Invitrogen PA5-29607 | Goat anti-rabbit HRP<br>1:1500 | Opal 540 1:1,000 |
| **K14 1:1000**<br>Invitrogen MA5-11599 | Goat anti-mouse HRP<br>1:1500 | Opal 520 1:150 |
| **Vimentin 1:300**<br>CST #5741 | Goat anti-rabbit HRP<br>1:1500 | Opal 690 1:150 |
| **E-cadherin 1:500**<br>BD #610182 | Goat anti-mouse HRP<br>1:1500 | Opal 650 1:500 |
| **ZEB1 1:1000**<br>Invitrogen PA5-82982 | Goat anti-rabbit HRP<br>1:1500 | Opal 570 1:600 |

#### Image Analysis

Whole slide scans were captured at 10x with the PerkinElmer Vectra3 Slide Scanner and ~50 regions of interest (ROIs) per tumor chosen manually with PhenoChart (PerkinElmer). ROIs were imaged at 20x resolution and imported into InForm analysis software (PerkinElmer). Spectral unmixing single-stains and background fluorescence slides were generated from the Parental tumor according to the OPAL Assay Development Guide. ROIs were spectrally unmixed and assigned colors and exported as composite images (Figure 4A). Tissue (Trainable to 98% accuracy) and cell segmentation were performed (Nuclear compartment – DAPI, Cytoplasm – Vimentin & KRT8, Membrane – E-cadherin) and cells were phenotyped based on expression of one or multiple markers (E-cadherin only, KRT8 & E-cadherin, KRT14 only, Triple positive (KRT8+E-cadherin+ Vimentin), Snail only, Vimentin only, and Vimentin+ZEB1) and validated by marker distribution (Supplemental Figure 5A&B). Entire Cell Mean Fluorescent units extracted for each marker and normalized as a percentile of maximum and minimum fluorescence across all cells in all images.

#### Heterogeneity and EMT Scores

##### Approach 1

Heterogeneity scores were generated using penalized logistic regression based on entropies of mean marker cell expressions to identify markers and cellular compartments (Nucleus, Cytoplasm, and Membrane) that contributed most to the variability in the ranked tumor images (Supplemental Figure 5C). In total 134 entropy-based features extracted, and 13 of them were selected by Recursive Feature Elimination (RFECV)^63^ as most relevant. Logistic regression classified sample heterogeneity into levels mid, low and high. Ground truths were determined from the rubric: low (one major cell trait with up to one minor trait), mid (two major cell traits with up to three minor traits), and high (three or more major cell traits present with two or more minor traits). These were used to train and validate the algorithm using 70% training and 30% test images (n=409) in a five-fold cross validation.

##### Approach 2

Nearest Neighbor analysis was conducted with the scikit-learn python package^45^ using cell phenotypes determined from InForm. Similar feature selection methods were applied to nearest neighbors, with 26 out of 49 features selected.

##### Approach 3

A hybrid approach used combined the 134 and 49 features from approach 1 and 2 and selected 18 out of 183 features.

To generate the EMT score, the seven derived phenotypes were weighted from epithelial to mesenchymal (E-cadherin only −3, KRT8 & E-cadherin −2, KRT14 only-1, Triple positive +1, Snail only +2, Vimentin only +3, and Vimentin & ZEB1 +4) and applied to a multivariate logistic regression.

### Raw Data

Raw data from RNA-seq and ATAC-seq experiments that support the findings in this study are currently being deposited in GEO. Accession codes will be provided once the submission is complete.

### Research Animals

All animal experiments were carried out under ethical regulations approved by the Dartmouth College IACUC

## Acknowledgements

We thank Jennifer Fields, Rebecca O’Meara, Dr. Radu V. Stan, Dr. Walburga Croteau and Dr. Fred W. Kolling IV for technical assistance; Molecular Biology Shared Resource at the Norris Cotton Center for assistance with RNA-Seq and ATAC-Seq assays; Dartlab (flow cytometry), microscopy and pathology shared resources at the Norris Cotton Cancer Center. Research reported in this publication was supported by an Institutional Development Award (IDeA) from the National Institute of General Medical Sciences of the National Institutes of Health (under grant number P20GM104416), a Prouty Pilot Grant from Friends of the Norris Cotton Cancer Center, funding from The Elmer R. Pfefferkorn & Allan U. Munck Education and Research Fund at the Geisel School of Medicine at Dartmouth and an NCI Cancer Center Support Grant (5P30CA023108-40). This work was supported by a Ramanujan Fellowship (SB/S2/RJN-049/2018) awarded by Science and Engineering Research Board (SERB), Department of Science and Technology (DST), Government of India (to M.K.J.) and funding from the NIH 5R00CA201574-05 (to D.R.P.).

## Author contributions

This project was conceived by M.S.B, N.B.O and D.R.P; N.B.O and M.S.B created the single cell EMT clones, which was the starting point for the project. M.S.B carried out all experiments except for generation of the entropy-based model for deriving heterogeneity scores and EMT scores, which was carried out by B.A and S.H, RNA-Seq and ATAC-Seq data analysis carried out by O.M.W and the EMT scoring by P.C and M.K.J. The manuscript was written by M.S.B and D.R.P with input and edits from all co-authors.

