## Supplemental materials and figures for "Dynamic plasticity within the EMT spectrum, rather than static mesenchymal traits, drives tumor heterogeneity and metastatic progression of breast cancers"

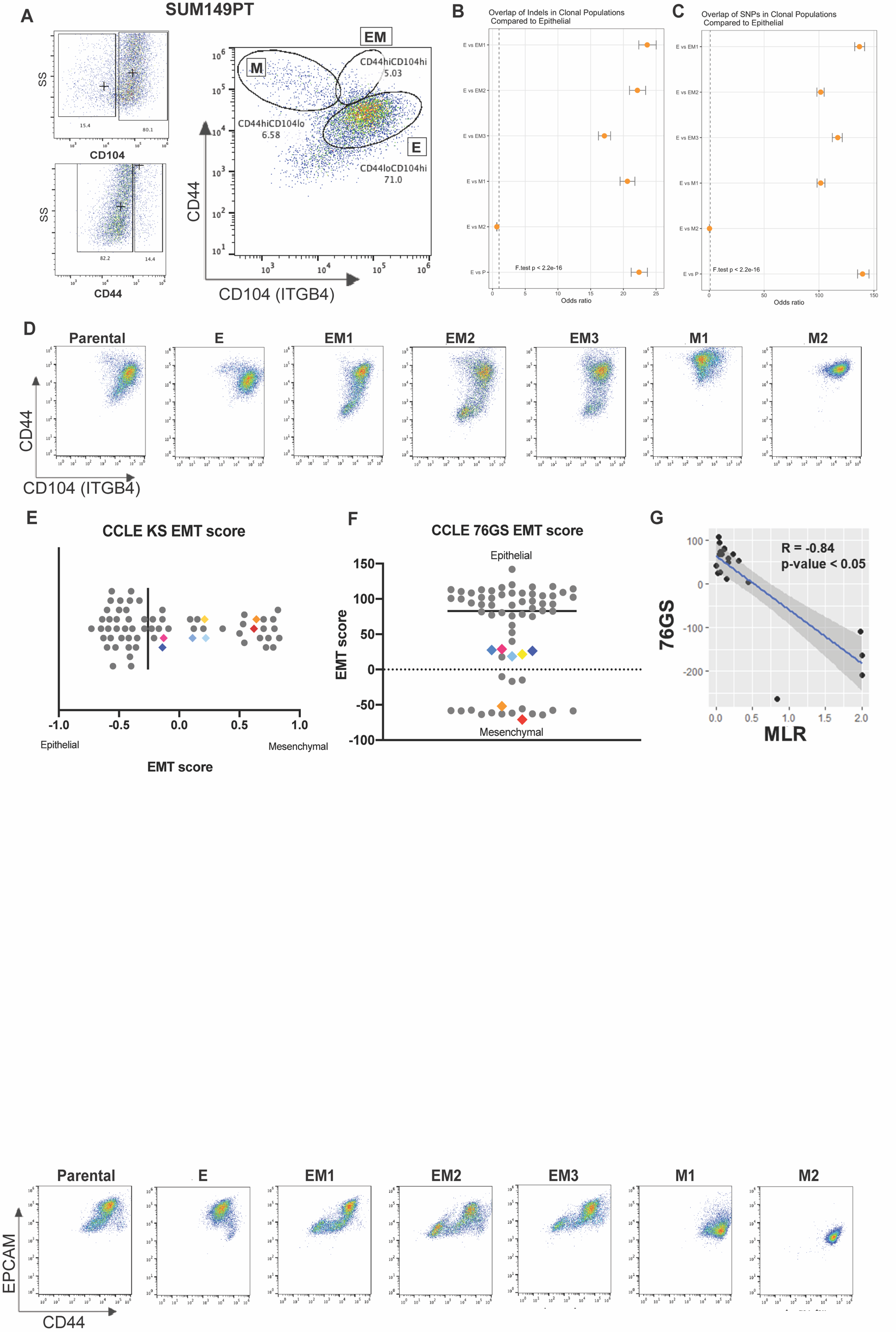

**Supplemental Figure 1:** **Heterogeneous cell line SUM149PT contains multiple distinct EMT states that can be isolated as single cell clones**

A) Fluorescence Assisted Cell Sorting (FACS) cytometry gating strategy to segregate individual EMT states from the heterogeneous breast cancer cell line SUM149PT, including epithelial (CD104^hi^CD44^lo^), intermediate (CD104^hi^CD44^hi^), and mesenchymal (CD104^lo^CD44^hi^) to generate clonal cell lines. B) Enrichment of clone specific variants (InDels and C) SNPs) among variants found in E. Fisher’s Exact Test was used to calculate the odds ratios and 95% confidence intervals shown on forest plots. Higher odds ratios indicate an increased odds that a variant will be shared between a given clone and E, suggesting relatively more genetic similarity. D) FACS plots of EMT clones stained with CD44 and CD104. E) EMT signature of EMT clones and parental line generated from the KS and F) 76GS methods of gene scoring plotted among other breast cancer cell lines from the Cancer Cell Line Encyclopedia (CCLE). G) Linear correlation between multiple EMT scoring metrics in the EMT clones.

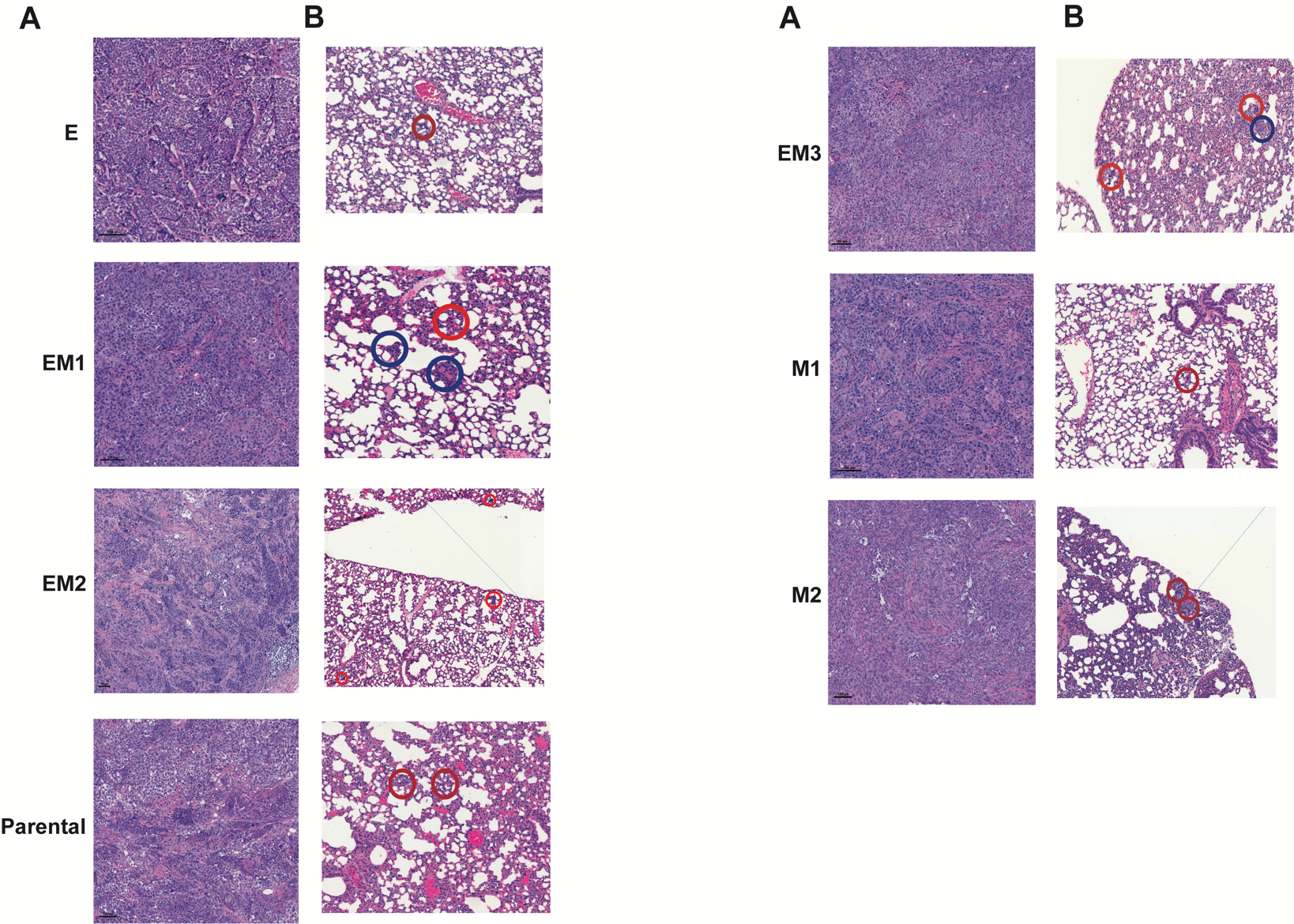

**Supplemental Figure 2: Histopathology of EMT clone tumors and staining of lungs to enumerate metastases**

A) Primary tumor stained with Hematoxylin & Eosin. B) Magnification of example lung metastases stained with Hematoxylin & Eosin. Micro- (red) and macro- (blue) metastases outlined

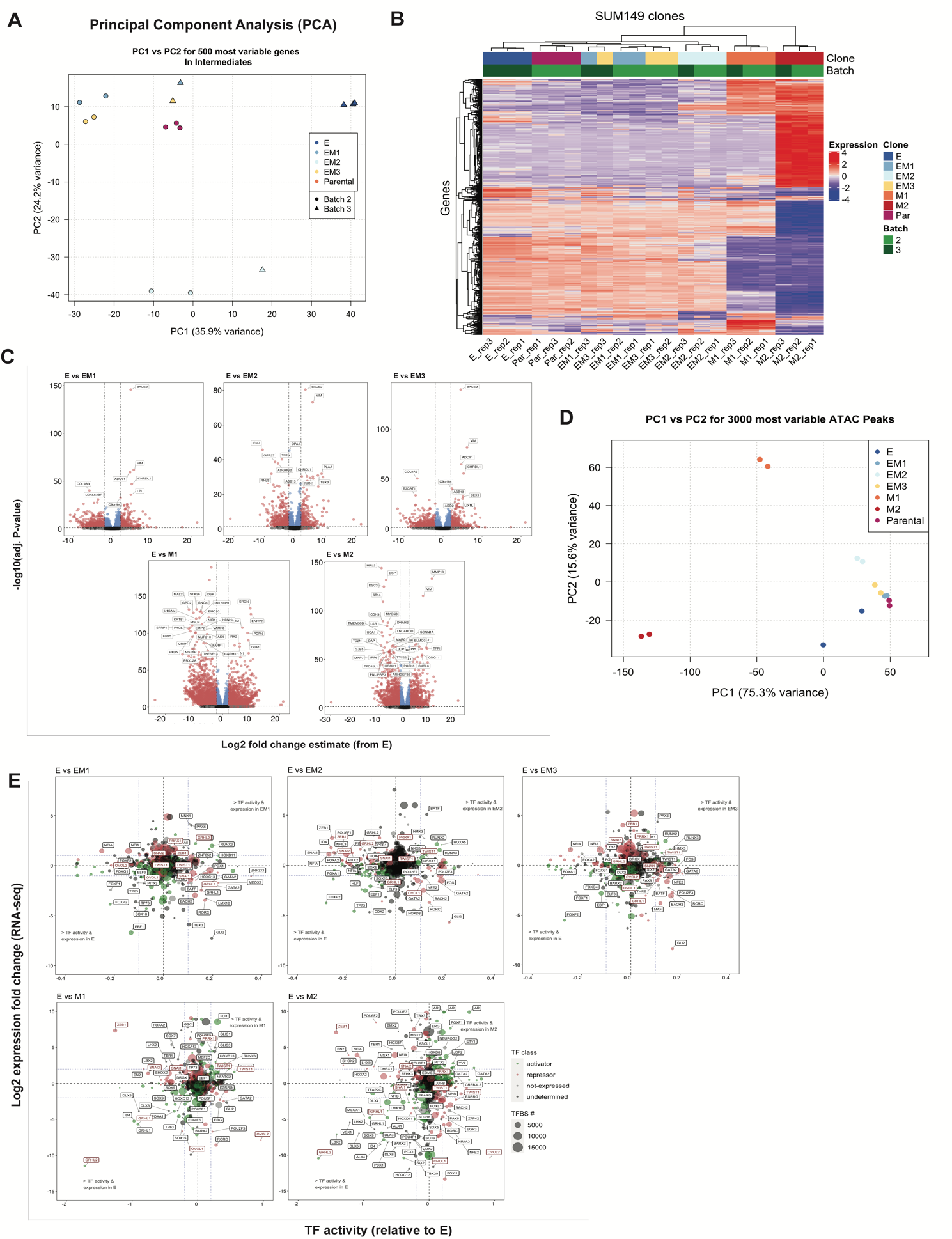

**Supplemental Figure 3: Identification of intermediate-stabilizing transcription factors by multi-omics analysis**

A) Principal Component Analysis (PCA) of the 500 most variably expressed genes across epithelial, intermediate EMT, and SUM149 Parental line taken from RNA-sequencing. Batch refers to sequencing run, normalized for batch effect. B) Unsupervised hierarchical clustering of the 500 most variably expressed genes across all clones. C) Volcano plots of showing differential expression analysis results from comparing each EMT clone in relation to clone E (reference group). Log_2_ fold change (LFC) from clone E on x-axis, -log_10_ adjusted p-value (FDR) on y-axis. Individual genes are color coded as follows: red, genes with adjusted p-value <0.05 and absolute LFC >2; blue, genes with adjusted p-value >0.05 & absolute LFC <2; gray, adjusted p-value <0.05 and absolute LFC >2; black, adjusted p-value >0.05 and absolute LFC >2. Select genes with most significant adjusted p-values and largest LFC are labelled. D) UpSet plot of all unique and shared differentially expressed genes in each clone compared to clone E. This shows the unique transcriptional landscape of each clone is more dominant than shared gene signatures between clones. E) Principal Component Analysis (PCA) of the 500 most variably expressed peak regions across epithelial, intermediate EMT, and SUM149 Parental line taken from ATAC-sequencing. F) Advanced volcano plot of highly significant transcription factors, highlighting the canonical EMT and MET driving transcription factors, relative to Epithelial determined by diffTF from ATAC-seq along the X-axis (label cutoffs at 0.1, -0.1 TF activity), plotted against Log2 fold gene expression values of transcription factors on the y-axis (label cutoffs at 1, -1 log2 fold). Transcription Factor classification, determined by transcription factor expression, shown in bubble color, and number of transcription factor binding sites (motifs) used to determine TF activity plotted as bubble size.

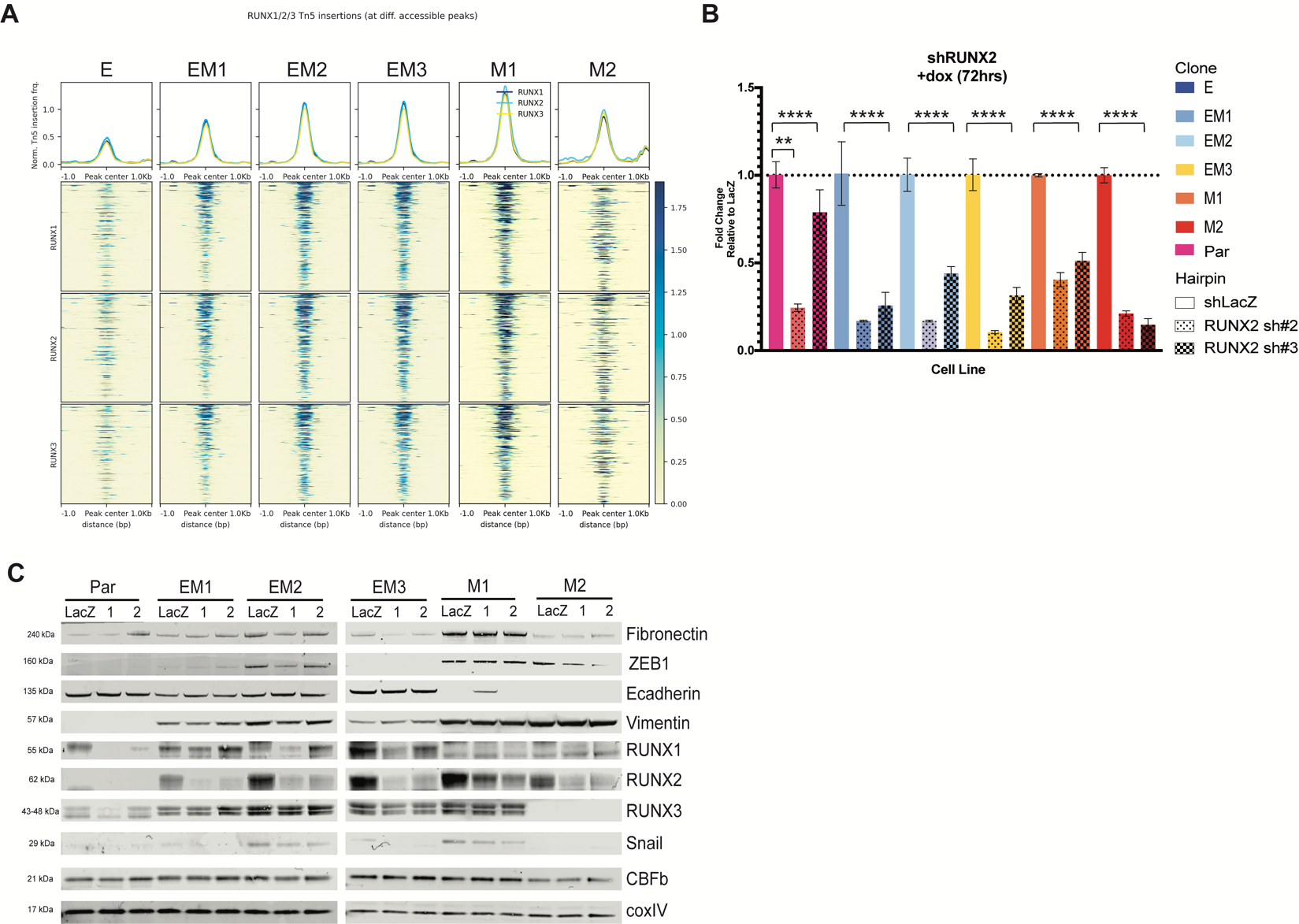

**Supplemental Figure 4: RUNX2 plays a stabilizing role in the intermediate state.**

A) ATAC-seq peak accessibility measured as counts per million (CPM) normalized Tn5 insertions surrounding RUNX1, RUNX2, and RUNX3 TF motifs for each clone. B) Confirmation of RUNX2 knockdown by inducible hairpin at 72 hours post-doxycycline with RT qPCR (** = 0.002, ****<0.0001) and C) Western blot.

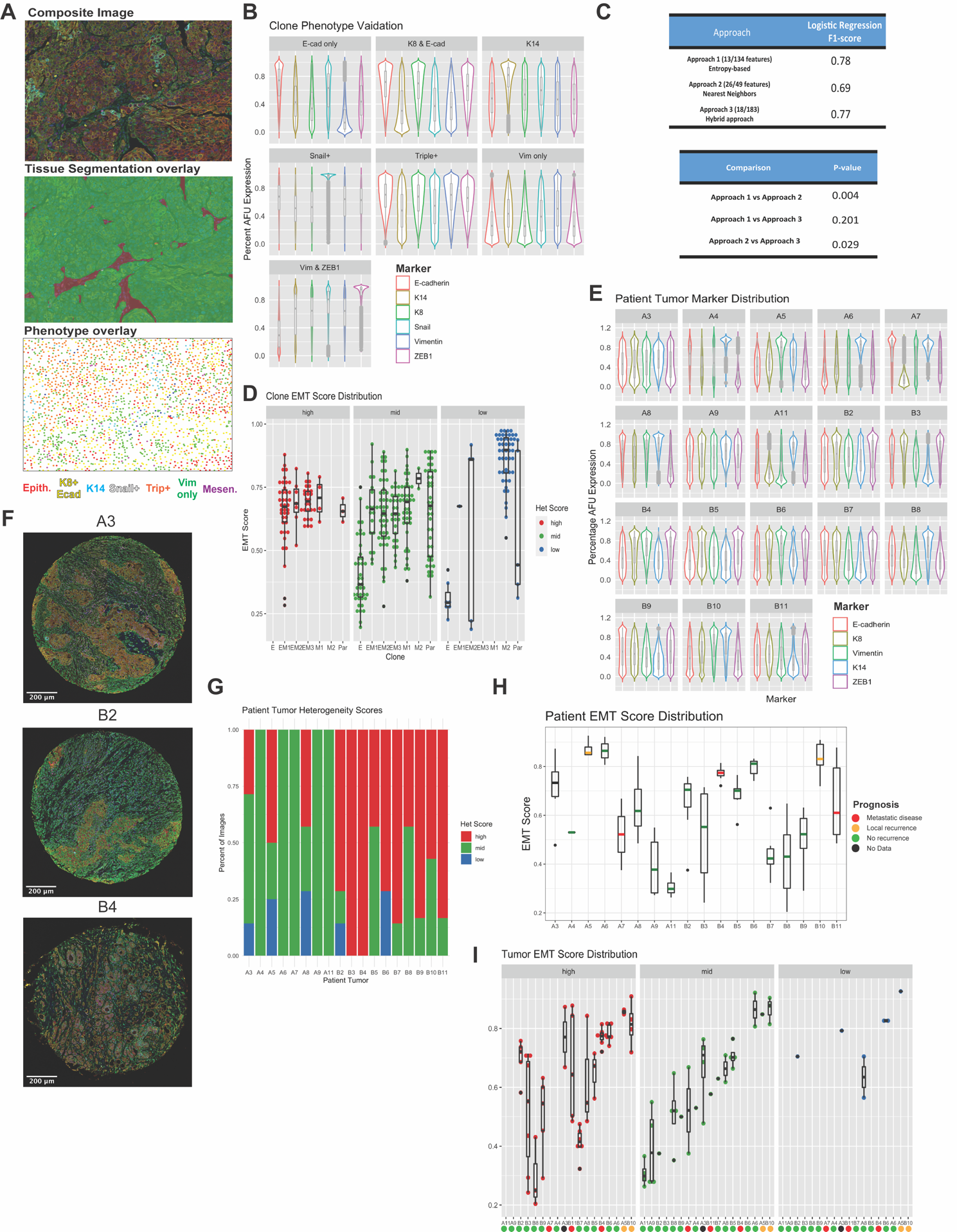

**Supplemental Figure 5: Multiplexed staining of tumors reveals shared phenotypes also found in patient tumors**

A) Outline of image analysis including Composite false colored image, trainable tissue segmentation, and trainable single cell phenotyping. B) Validation of EMT phenotype selection. Normalized EMT marker distribution (Percent AFU Expression) for each phenotype shows enrichment for markers described in the phenotypes (i.e., K14 phenotyped cells are enriched for high expression of K14). C) Entropy-based logistic regression scoring for three approaches to determining heterogeneity. (1) an entropy-based approach using mean marker expression of cells per image, determined by 13 out of 134 entropy-based feature interactions (2) a nearest neighbor analysis approach using cell phenotypes as determined in B), determined by 26 out of 49 feature interactions and (3) a hybrid approach combining approach 1 and 2, determined by 18 of 183 features combined from approach 1 and 2. D) EMT clone correlation between EMT score and heterogeneity score. Each dot represents one image from an EMT clone-derived tumor. E) EMT percentile normalized marker distribution of all staining markers across 18 patient tumors stained as in (A). F) Examples of patient tumors stained with multiplexed EMT markers showing complementary phenotypes and architecture to EMT model clones. G) Patient tumor heterogeneity scores determined by Approach 1 from (C) after multiplexed staining for EMT markers. H) Patient tumor EMT score distribution generated from weighted multivariable logistic regression of the phenotypes present in each tumor. Median line color indicates patient prognosis. I) Correlation of EMT score and Heterogeneity score between patient tumors. Each dot represents an image. Colored dots at the x-axis indicate patient prognosis.
